# A multivariate pattern metric of individualized hemispheric functional asymmetry

**DOI:** 10.64898/2025.12.03.692004

**Authors:** Qiuhui Bi, Chenxi Zhao, Xi-Nian Zuo, Shaoling Peng, Bo Sun, Gaolang Gong

## Abstract

Functional asymmetry is a fundamental feature of human brain organization, yet existing measures of functional connectivity asymmetry rely mainly on univariate indices that miss distributed pattern structure. We introduce the Pattern Dissimilarity of Hemispheric Functional Connectivity (PDHC), a multivariate metric that quantifies the pattern-level distance between each individual’s left- and right-hemisphere connectivity architecture. In over one thousand adults, PDHC showed high test-retest reliability, cross-atlas robustness, and strong individual specificity. Network analyses indicated that hemispheric similarity is anchored in conserved sensorimotor and subcortical systems, whereas higher-order networks drive divergence. Developmental data from infancy through early adulthood revealed a characteristic trajectory: dissimilarity declines sharply early in life and increases modestly thereafter. Twin and Turner syndrome samples further demonstrated moderate heritability and sensitivity to X-chromosome dosage. PDHC thus provides a reliable and individualized metric of hemispheric functional architecture, offering a scalable tool for probing lateralization across development, genetics, cognition, and clinical conditions.

## Introduction

Despite their shared evolutionary and developmental origins, the two cerebral hemispheres are not functionally equivalent. Substantial evidence demonstrates hemispheric lateralization and asymmetry across diverse domains such as language, attention, and emotion (Güntürkün & Ocklenburg, 2017; Ocklenburg & Güntürkün, 2024). Functional magnetic resonance imaging (fMRI) has played a central role in delineating these asymmetries across task and resting states. In particular, resting-state fMRI (rs-fMRI) studies have revealed consistent hemispheric differences in functional connectivity (FC)— the temporal synchrony of spontaneous neural activity across brain regions (Gotts et al., 2013; Joliot et al., 2016; Liang et al., 2021; Perez et al., 2023; Tzourio-Mazoyer et al., 2016). However, these findings predominantly relied on univariate analyses, in which each FC link is treated as an independent feature, thereby obscuring the distributed and systems-level nature of hemispheric functional divergence (Averbeck et al., 2006; Karolis et al., 2019; Zhu et al., 2022) and limiting our understanding of asymmetry as an emergent property of large-scale functional architectures.

While single connections can capture localized differences, the human brain operates as an integrated system whose functional architecture is best described by distributed activity patterns (Bertolero et al., 2015; Petersen & Sporns, 2015; Smallwood et al., 2021). Multivariate approaches—such as multi-voxel pattern analysis (MVPA)—have revolutionized the study of neural representation by decoding population-level information embedded in spatially distributed fMRI signals (Haxby et al., 2001; Norman et al., 2006; Tong & Pratte, 2012). Extending this concept to the connectivity domain, recent work has shown that whole-brain FC patterns carry unique individual signatures, can distinguish between cognitive states, and predict behavioral traits or brain maturity (Cui & Gong, 2018; Finn et al., 2015; Gonzalez-Castillo et al., 2015; Jo et al., 2021; Shirer et al., 2012). Despite their widespread success, multivariate frameworks have seldom been employed to quantify hemispheric functional asymmetry, leaving unresolved a central question in systems neuroscience: how different, as complete network organizations, are the two cerebral hemispheres, and what principles govern their pattern-level divergence?

The left and right hemispheres form two largely autonomous yet interacting systems. Evidence from callosotomy and corpus callosum agenesis shows that intra-hemispheric FC patterns remain remarkably stable even when interhemispheric communication is severely disrupted (Tyszka et al., 2011; Uddin et al., 2008). Similarly, hemispherectomy patients retain well-organized connectivity architectures within the preserved hemisphere that closely resemble those in typical individuals (Kliemann et al., 2019, 2021). These observations indicate that each hemisphere harbors an internally coherent, self-sufficient functional organization. Recognizing this autonomy raises a fundamental question: to what extent do the hemispheres instantiate distinct and diverging network-level architectures, beyond the asymmetry revealed by any individual connection or region? Quantifying such pattern-level dissimilarity offers a principled way to characterize functional asymmetry beyond conventional laterality indices. Yet no existing framework enables a direct, multivariate comparison of the two hemispheres as high-dimensional representationalsystems, representing a conceptual and methodological gap in the study of hemispheric specialization.

Here, we introduce a novel multivariate metric—Pattern Dissimilarity of Hemispheric Functional Connectivity (PDHC)—that quantifies the dissimilarity between whole-hemisphere FC patterns. Conceptually, PDHC captures the pattern-level distance between the two hemispheric connectivity architectures, providing an explicit representational-space formulation of hemispheric specialization that extends traditional laterality indices into a high-dimensional, systems-level framework. This formulation enables hemispheric divergence to be characterized as an intrinsic architectural property of large-scale functional networks, independent of specific cognitive or behavioral measures. Empirically, we demonstrate that PDHC is (i) reproducible across acquisition sessions and parcellation schemes; (ii) developmentally dynamic, showing systematic changes in three independent cohorts; and (iii) genetically influenced, as revealed by twin and Turner syndrome data. Collectively, these findings establish PDHC as a robust and interpretable marker of hemispheric functional specialization, offering a unifying framework for quantifying how hemispheric pattern-level divergence shape human brain organization across individuals, developmental stages, and genetic backgrounds. By providing the first direct multivariate assessment of hemispheric functional architectures, PDHC addresses a longstanding question in systems neuroscience: how bilateral brains are organized, maintained, and varied in their intrinsic large-scale architecture.

## Results

### High reliability and robustness of PDHC

Resting-state fMRI data from 1,033 HCP young adults were analyzed (Table S1). PDHC was defined as the Pearson-correlation distance between left- and right-hemispheric functional-connectivity (FC) matrices, quantifying whole-brain functional asymmetry (Fig. 1A). PDHC exhibited high reproducibility across both short- and long-term intervals. In 960 participants scanned twice approximately one day apart, REST1 and REST2 distributions overlapped substantially (K–S Z = 1.0, p > 0.20), and session-wise correlations were strong (r = 0.63–0.67; ICC = 0.63–0.67) (Fig. 1B). Over longer intervals of up to 11 months, reliability remained high (r = 0.62–0.65; ICC = 0.63–0.65), demonstrating that PDHC is stable on both short- and extended timescales (Fig. 1C).

**Figure 1.**
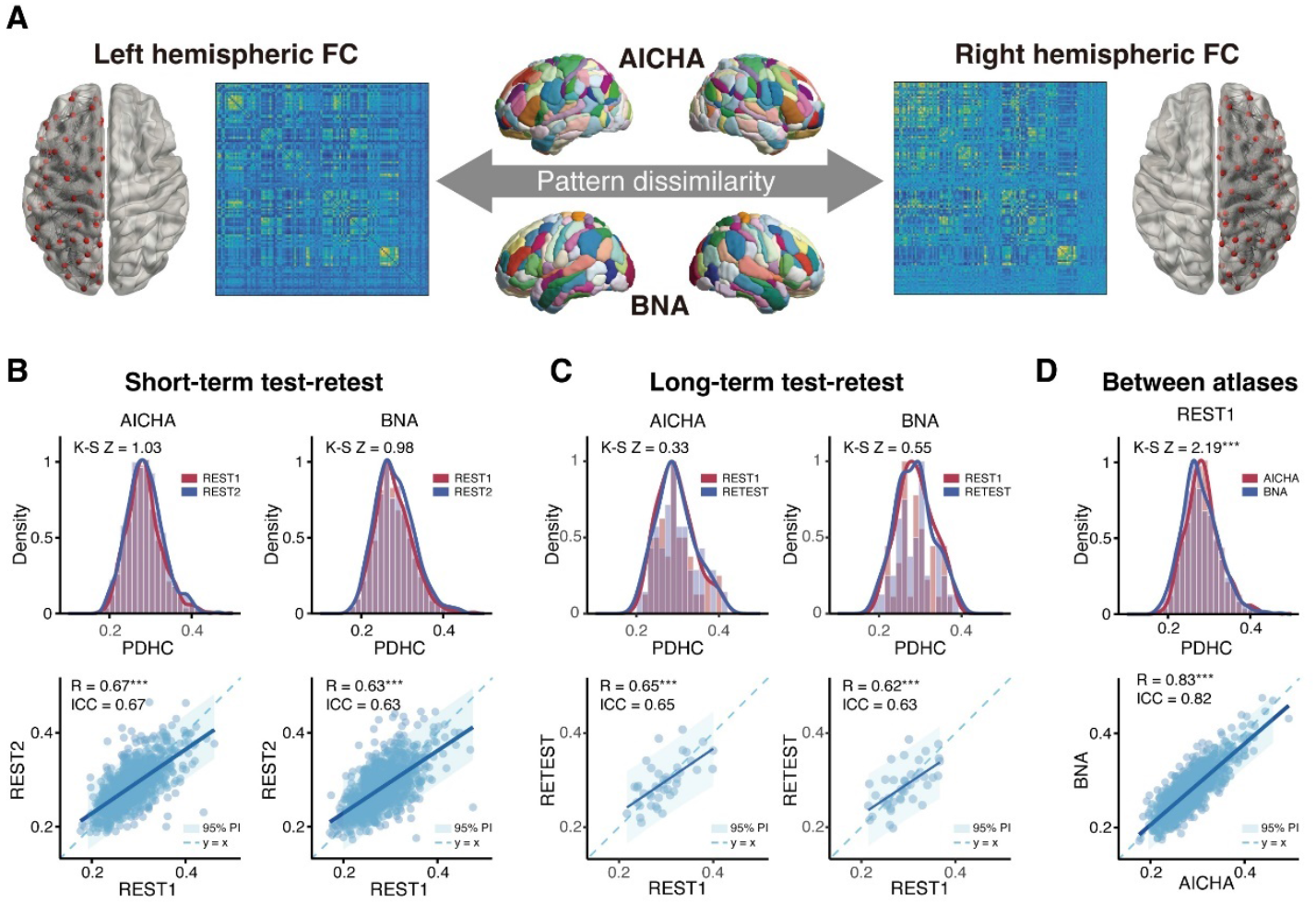
Reliability and cross-atlas robustness of the Pattern Dissimilarity of Hemispheric Functional Connectivity (PDHC). (A) Schematic of PDHC computation. For each participant, functional-connectivity (FC) matrices were constructed separately for the left and right hemispheres using the AICHA and BNA atlases. PDHC was defined as the Pearson-correlation distance between the two hemispheric FC patterns, quantifying the pattern-level dissimilarity of large-scale hemispheric organization. (B) Short-term (one-day) test–retest reproducibility. PDHC distributions showed strong overlap between REST1 and REST2 in both atlases, and session-wise correlations (r = 0.63–0.67; ICC = 0.63–0.67) indicated high short-term reproducibility. (C) Long-term test–retest reliability. PDHC remained stable over months, with robust REST1–RETEST correlations (r = 0.62–0.65; ICC = 0.63–0.65) and largely overlapping empirical distributions. (D) Cross-atlas correspondence. Although mild shifts were detected by K–S tests, inter-atlas correlations were high (r = 0.83; ICC = 0.82), demonstrating that individual differences in PDHC are robust to parcellation scheme. Abbreviations: PDHC, Pattern Dissimilarity of Hemispheric Functional Connectivity; FC, functional connectivity; AICHA, Atlas of Intrinsic Connectivity of Homotopic Areas; BNA, Brainnetome Atlas; PI, prediction interval.

PDHC values were also highly concordant across parcellation schemes (r = 0.83; ICC = 0.82), with only minor distributional shifts (Fig. 1D), and were robust to preprocessing variations, including omission of global-signal regression (Fig. S1). Associations with motion, scan length, and other nuisance variables were minimal (< 5% variance explained). Reanalysis of an unrelated HCP subset reproduced these patterns (r = 0.59–0.80; ICC = 0.59–0.80), indicating independence from familial structure. Moreover, short-term reproducibility was comparable across atlases, and long-term stability persisted after accounting for scanner drift and physiological variability, supporting the trait-like nature of PDHC.

### Individual specificity and distinctiveness of PDHC

To assess whether PDHC captures individualized hemispheric organization, we compared intra-subject PDHC values with inter-subject baselines derived from randomly paired individuals. One-way ANOVA revealed large effects (all p < 10^− 22^), and post-hoc tests confirmed that intra-subject PDHC values were consistently lower than inter-subject ipsilateral and contralateral dissimilarities (all t < –90, all p < 0.001; Fig. 2A). These effects were replicated across both AICHA and BNA atlases and across REST1 and REST2.

**Figure 2.**
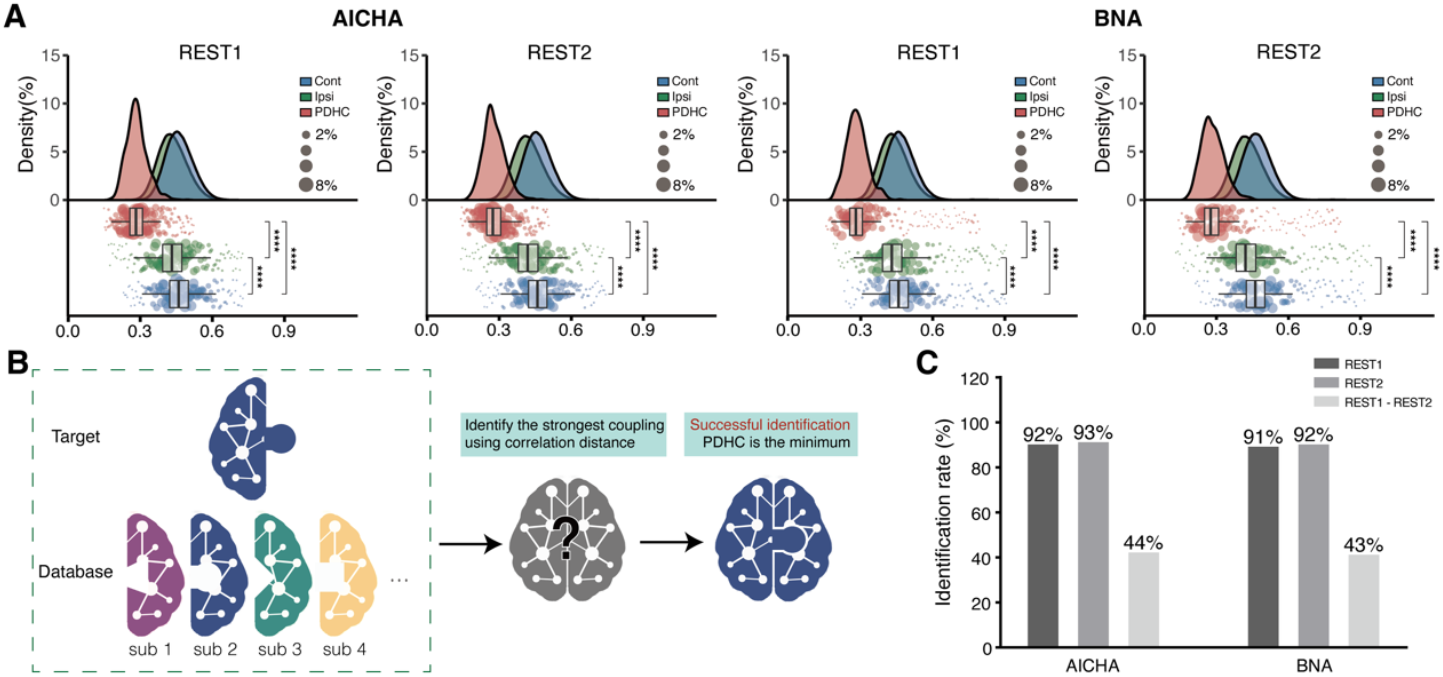
PDHC captures individualized hemispheric dissimilarity. (A) Within-versus between-subject hemispheric dissimilarity. Across both AICHA and BNA atlases, intra-subject PDHC values were consistently lower than between-subject baselines generated from ipsilateral and contralateral pairings (all p< 0.001). Upper panels show kernel density estimates; lower panels show boxplots with density-weighted scatter points (point size proportional to sample density). Boxplots depict the interquartile range (IQR) with median lines; whiskers extend to 1.5×IQR. (B) Subject-identification framework. For each target hemisphere, correlation distances to all hemispheric connectomes in the database were computed, and identification was deemed successful when the within-subject PDHC was the minimum across all pairwise distances. (C) Individual identification accuracy. Within-session identification was high (91–93%), whereas cross-session identification (REST1 → REST2) was moderate (43–44%), reflecting stable yet session-dependent individuality in hemispheric FC patterns. Abbreviations: PDHC, Pattern Dissimilarity of Hemispheric Functional Connectivity; FC, functional connectivity; AICHA, Atlas of Intrinsic Connectivity of Homotopic Areas; BNA, Brainnetome Atlas.

To further test subject specificity, we implemented a subject-identification framework (Fig. 2B). A participant was considered successfully identified when their intra-subject PDHC was the minimum across all distances. Identification accuracy was high within sessions (92–93%) and moderate across sessions (43–44%) (Fig. 2C). When restricting analyses tounrelated participants, accuracies increased further (within-session: 97–99%; cross-session: 68–72%), indicating that PDHC reflects stable person-specific hemispheric organization rather than family similarity or transient state effects. Thus, PDHC encodes a reproducible individual fingerprint of hemispheric organization, highlighting enduring left–right pattern-level differentiation as an intrinsic neural trait.

### Network architecture and contributing systems of PDHC

To determine which functional systems contribute most to hemispheric dissimilarity, we decomposed PDHC into edge-wise standardized contributions, z-scored relative to a permutation-based null. Because PDHC quantifies hemispheric dissimilarity, negative edge-wise contributions correspond to features that reduce left–right dissimilarity. These contributions were projected onto both Mesulam’s hierarchical parcellation and Yeo’s 7-network taxonomy, allowing system-level inference across complementary organizational schemes (Table S2; Fig. S3). Enrichment analysis identified network pairs disproportionately represented among edges with the largest negative contributions. Overrepresentation was quantified using the rich factor (RF) and evaluated via 5,000 permutation tests with FDR correction, thereby identifying systems containing a greater-than-chance density of dissimilarity-reducing edges.

Nearly all edges contributed negatively (Fig. S4), indicating that most functional connections act to reduce hemispheric dissimilarity at the whole-connectome level. As shown in Fig. 3, in Mesulam’s hierarchy, Primary–Primary edges showed the strongest enrichment (pFDR < 0.001), followed by Primary–Limbic/Subcortical and Primary– Unimodal pairs (Table S3). In Yeo’s taxonomy, VIS–SMN and SMN–SMN pairs were most enriched (pFDR < 0.005), with further contributions from VIS–VAN and SMN– Subcortical pairs (Table S4). These convergent patterns indicate that primary sensorimotor and subcortical systems provide the dominant connectivity architecture that minimizes hemispheric dissimilarity, consistent with their conserved and bilaterally organized functional roles. Enrichment patterns were stable across thresholds (3–20%) and consistent between the AICHA and BNA parcellations (Fig. S5).

**Figure 3.**
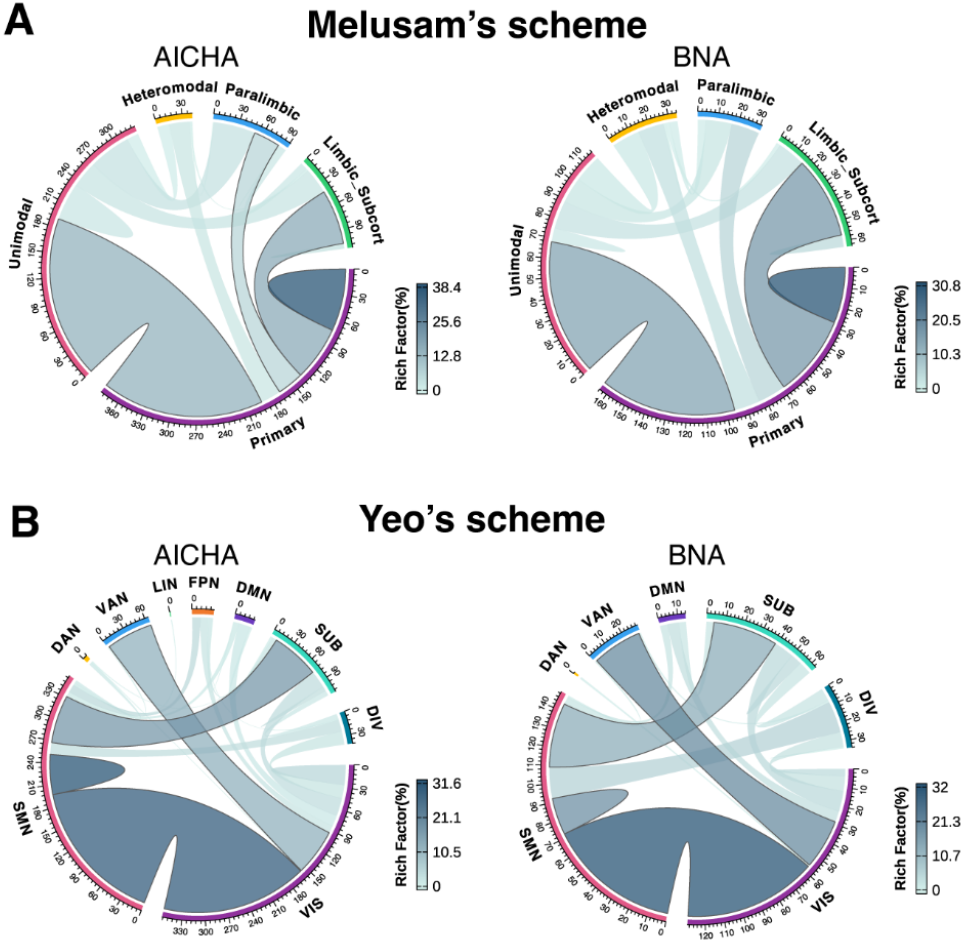
Network-level enrichment of edges contributing to hemispheric dissimilarity as quantified by PDHC. Chord diagrams show enrichment of edges with the strongest negative PDHC contributions (i.e., edges that reduce left–right dissimilarity), thresholded at the top 3% using REST1 data for both AICHA and BNA atlases. Upper panels display enrichment within Mesulam’s hierarchical organization, and lower panels map results onto Yeo’s 7-network system. Line color encodes the rich factor (RF), line width reflects the number of edges exceeding the threshold, and bold segment borders denote FDR-corrected significance (pFDR < 0.05). Axes represent unnormalized network-level edge counts. Abbreviations: PDHC, Pattern Dissimilarity of Hemispheric Functional Connectivity; RF, rich factor; FDR, false discovery rate; VIS, visual; SMN, sensorimotor; VAN, ventral attention; DAN, dorsal attention; LN, limbic network; FPN, frontoparietal network; DMN, default mode network; Subc, subcortical.

### Developmental dynamics of PDHC from infancy to adulthood

To characterize the maturation of hemispheric functional asymmetry, we evaluated PDHC in three large developmental cohorts: neonates (dHCP), children/adolescents (devCCNP), and young adults (HCP). Linear mixed-effects models revealed a pronounced decrease of PDHC with postmenstrual age (PMA) in neonates (t = -17–-16, p < 10^-43^; Fig. 4A), even after adjusting for delivery type, birth weight, sex, head circumference, and motion. The trend persisted in full-term infants (t = -8.5–-8.0, p < 10^-13^) and low-motion infants (r = - 0.81, p < 10^-50^).

**Figure 4.**
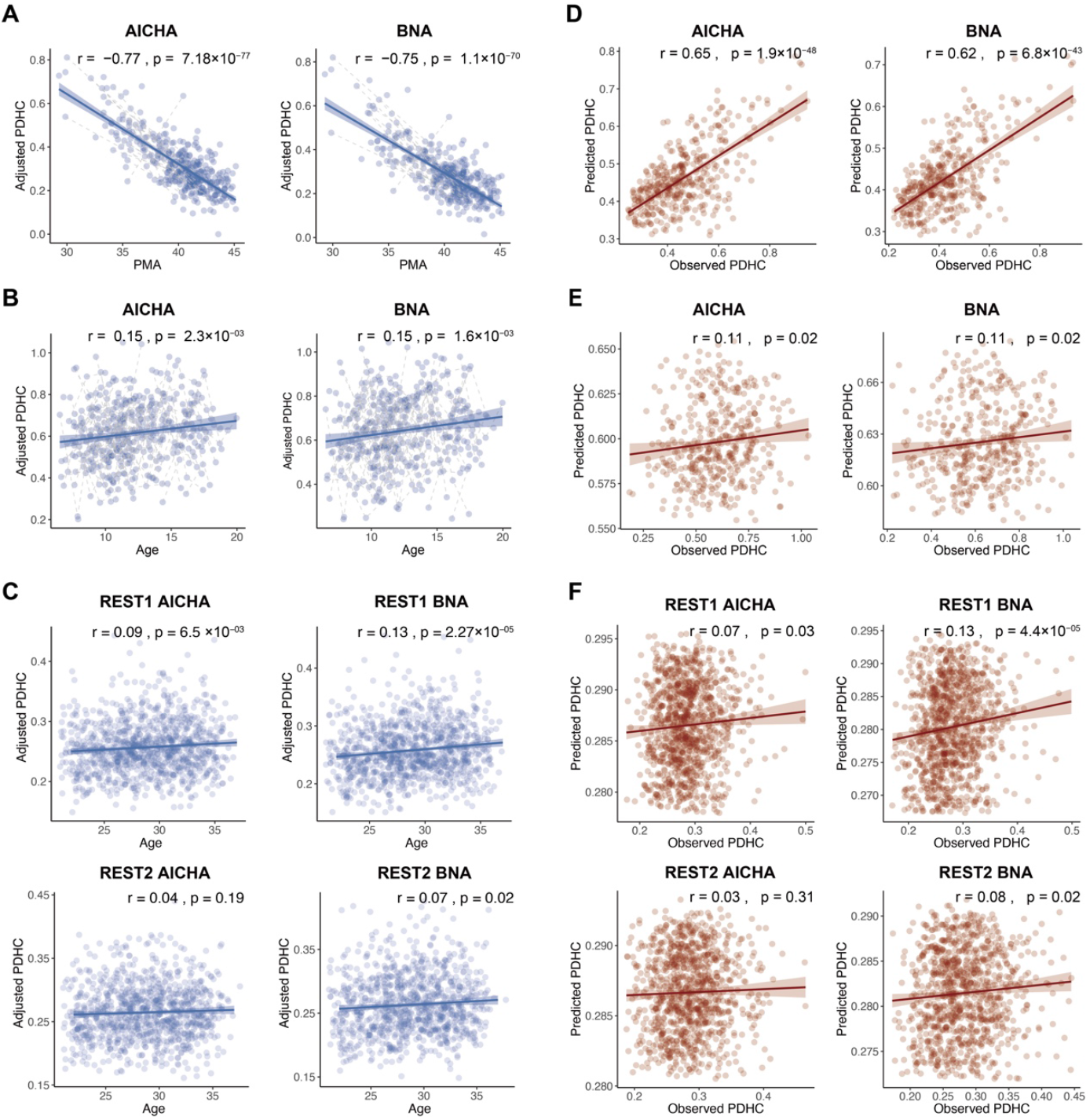
Developmental trajectory and age predictability of PDHC from infancy to adulthood. (A) In neonates, PDHC decreased sharply with postmenstrual age (PMA) in both AICHA and BNA atlases, reflecting rapid early differentiation of hemispheric architecture. (B) From childhood to adolescence, PDHC showed modest age-related increases, suggesting continued maturation of hemispheric specialization. (C) In early adulthood, PDHC showed weak but detectable positive age trends, representing stabilization rather than continued differentiation of hemispheric organization. (D–F) Age prediction using leave-one-out cross-validation (LOOCV). Predicted and observed PDHC values were highly correlated in neonates, significantly correlated in children/adolescents, and minimally correlated in adults, consistent with developmental changes in variability. Longitudinal datasets (dHCP, devCCNP) were analyzed using linear mixed-effects models, and the cross-sectional HCP dataset was analyzed using ordinary linear models. Abbreviations: PDHC, Pattern Dissimilarity of Hemispheric Functional Connectivity; PMA, postmenstrual age; AICHA, Atlas of Intrinsic Connectivity of Homotopic Areas; BNA, Brainnetome Atlas; LOOCV, leave-one-out cross-validation; dHCP, developing Human Connectome Project; devCCNP, developmental Chinese Color Nest Project; HCP, Human Connectome Project.

In the devCCNP cohort, PDHC showed a modest but significant increase with age (t = 2.7– 2.8, p = 0.005–0.007; Fig. 4B), although individual variability was substantial (r = 0.15– 0.20). Among HCP young adults, PDHC exhibited weak positive age associations (Fig. 4C). Leave-one-out cross-validation demonstrated that PDHC strongly predicted age in infants (r = 0.62–0.66, p < 10^-40^) but was far less predictive in adolescence and adulthood (r = 0.04–0.13; Fig. 4D–F). Analyses restricted to first sessions of dHCP and devCCNP confirmed these results (Fig. S2). Together, these findings delineate a developmental trajectory in which hemispheric dissimilarity decreases rapidly during late gestation and early infancy, increases modestly throughout childhood and adolescence, and stabilizes into early adulthood, consistent with progressive specialization and refinement of hemispheric functional architecture.

### Genetic basis and heritable influences on PDHC

To examine the biological origins of hemispheric functional asymmetry, we assessed the genetic influence on PDHC. Twin analyses in the HCP cohort showed higher intraclass correlations in monozygotic twins (ICC_MZ_ = 0.32–0.47) than in dizygotic twins (ICC_DZ_ = 0.05–0.16), yielding significant heritability estimates (h^2^ = 0.25–0.42, all p < 0.001; Fig. 5A). These effects were consistent across REST1 and REST2 and robust to covariate adjustment for motion and global signal.

**Figure 5.**
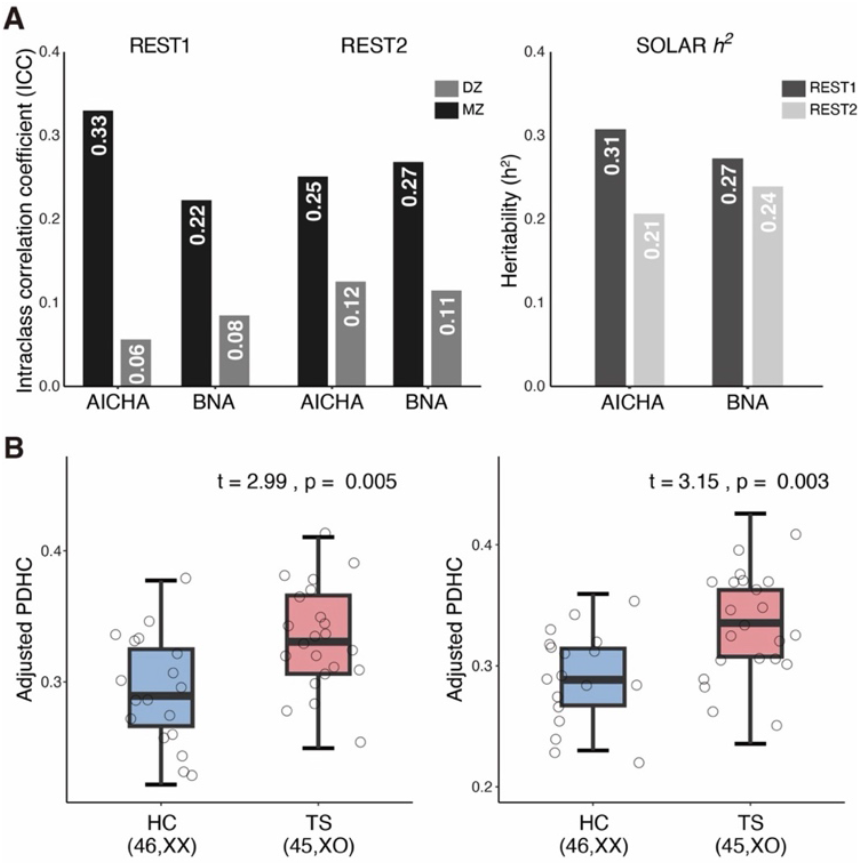
Genetic influences on hemispheric pattern asymmetry measured by PDHC. (A) Intraclass correlation coefficients (ICCs) showed greater within-pair similarity for monozygotic (MZ) versus dizygotic (DZ) twins across REST1 and REST2, and heritability estimates (h^2^) derived via SOLAR indicated moderate heritability in both AICHA and BNA. (B) Adjusted PDHC values were significantly higher in Turner syndrome (TS; 45,XO) compared with age-matched healthy controls (HC; 46,XX) across both parcellations (AICHA: t = 2.99, p = 0.005; BNA: t = 3.15, p = 0.003). Abbreviations: PDHC, Pattern Dissimilarity of Hemispheric Functional Connectivity; ICC, intraclass correlation coefficient; MZ, monozygotic twins; DZ, dizygotic twins; h^2^, heritability; TS, Turner syndrome; HC, healthy controls; AICHA, Atlas of Intrinsic Connectivity of Homotopic Areas; BNA, Brainnetome Atlas.

Complementary evidence came from Turner syndrome. Individuals with X-chromosome monosomy (45,XO) exhibited significantly elevated PDHC compared with age-matched healthy controls (HC; 46,XX) across both AICHA and BNA parcellations (p < 0.01; Fig. 5B). These results indicate that hemispheric pattern-level asymmetry is moderately heritable and sensitive to X-linked genetic dosage, supporting PDHC as a functional phenotype that bridges structural, developmental, and genetic influences on hemispheric organization.

## Discussion

This study introduces Pattern Dissimilarity of Hemispheric Functional Connectivity (PDHC), a multivariate framework for quantifying hemispheric asymmetry in functional connectivity patterns. Unlike conventional laterality indices that measure directional biases in individual connections (Gotts et al., 2013; Toga & Thompson, 2003), PDHC captures the magnitude of dissimilarity between distributed intrahemispheric connectivity architectures. By providing a non-directional, pattern-level measure, PDHC quantifies how distinct the two hemispheric functional architectures are, independent of which hemisphere shows greater activity or connectivity. In this way, PDHC characterizes hemispheric divergence in terms of its geometry rather than its sign, revealing a previously uncharacterized dimension of large-scale functional asymmetry. This multivariate perspective expands existing conceptualizations of hemispheric specialization by emphasizing distributed architectural differentiation rather than focal dominance, thereby offering a systems-level account of how functional specialization is embedded within the human connectome.

Traditional laterality metrics have been instrumental in mapping localized functional dominance, particularly in language or emotion networks (Hervé et al., 2013; Tzourio-Mazoyer et al., 2015). However, such univariate approaches treat each connection asindependent and collapse complex interactions into a scalar index. PDHC transcends these limitations by comparing entire hemispheric connectivity matrices, thereby capturing the distributed organization of hemispheric functional connectivity (Finn et al., 2015; Krienen et al., 2014; Yeo et al., 2014). Within this multivariate framework, functional asymmetry emerges as a global property of the connectome—a network-level configuration shaped by interactions among many connections rather than a few lateralized pairs. PDHC thus reframes hemispheric specialization as a property of functional geometry within shared structural constraints. This perspective aligns with contemporary theories emphasizing representational manifolds, population coding, and distributed computation as canonical modes of brain organization (Averbeck et al., 2006; Bassett & Sporns, 2017; Cunningham & Yu, 2014; Kriegeskorte, 2015), situating hemispheric asymmetry within these broader principles.

At the systems level, PDHC decomposition uncovered a hierarchical pattern of hemispheric dissimilarity reduction, with lower-order networks—sensorimotor, visual, and subcortical—dominating contributions that minimize pattern-level differences (Biswal et al., 2010; Stark et al., 2008). These findings suggest that large-scale hemispheric divergence is constrained by evolutionarily conserved bilateral systems that maintain high cross-hemispheric coordination to support perceptual and motor integration (Buckner & Krienen, 2013; D. Wang et al., 2014). In contrast, association cortices contributed minimally to dissimilarity reduction at rest, consistent with their flexible, state-dependent functional profiles (Cole et al., 2013; de Pasquale et al., 2012; Raichle, 2015). These patterns support a principle of hierarchically graded differentiation: bilateral coherence in lower-order networks provides a scaffold from which higher-order lateralized functions emerge. Importantly, this systems hierarchy offers a mechanistic backdrop for interpreting developmental effects, because early-maturing primary systems establish the bilateral scaffold upon which later-developing association cortices differentiate (Keller et al., 2023; Larsen et al., 2023; Sydnor et al., 2021). Because these findings are derived from resting-state data, future task-based analyses will be essential to determine how PDHC reorganizes during lateralized cognitive operations.

Across development, PDHC delineates a coherent trajectory of hemispheric functional asymmetry that aligns with major maturational shifts in large-scale network organization. Neonates exhibited high PDHC values that decreased sharply with age, indicating a rapid reduction in hemispheric dissimilarity as early functional circuits become increasingly coordinated (Cao et al., 2017; Emerson et al., 2016). During childhood and adolescence, PDHC showed modest age-related increases, consistent with the gradual emergence of more differentiated functional architectures as association networks mature (Menon, 2013; Power et al., 2010; J. Wang et al., 2020). In early adulthood, PDHC demonstrated only weak positive age trends, suggesting that the broad pattern of hemispheric organization has largely stabilized by this stage (Betzel et al., 2014; Edde et al., 2021). Together, these developmental changes reflect a transition from early bilateral integration toward increasingly structured hemispheric specialization, mirroring the developmental sequence in which primary systems mature early while association cortices undergo prolonged refinement (Buckner & Krienen, 2013; Larsen et al., 2023; Sydnor et al., 2021).

Genetic analyses further position PDHC within the biological architecture of hemispheric organization. Twin-based heritability and X-chromosome effects observed in Turner syndrome together indicate that hemispheric functional asymmetry reflects intertwined genetic and developmental influences (Güntürkün & Ocklenburg, 2017; Sha et al., 2021). The moderate heritability of PDHC suggests a polygenic basis that operates alongside environmental influences, positioning PDHC as a biologically grounded phenotype for investigating genetic contributions to hemispheric specialization. These results also highlight PDHC’s potential as an intermediate phenotype for imaging–genetics studies aimed at identifying molecular pathways underlying hemispheric differentiation.

Conceptually, PDHC complements directional laterality indices by quantifying how much the hemispheres differ rather than which side dominates. This distinction parallels the difference between measuring variance and measuring mean bias—both informative but capturing different organizational principles. The absence of directional sign is advantageous in systems where specialization reflects distributed reconfiguration rather than unilateral enhancement (Duncan et al., 2020; Westerhausen et al., 2014), allowing PDHC to provide a stable and generalizable index of asymmetry across domains with mixed or uncertain lateralization (Friederici & Alter, 2004; Kong et al., 2018; Yuan et al., 2021). In this way, PDHC overcomes a longstanding limitation of traditional laterality metrics, which often struggle to quantify asymmetries that are distributed, multidimensional, or non-monotonic.

Methodologically, PDHC’s multivariate nature offers a scalable platform for cross-modal extension. Its core principle—pattern dissimilarity between homologous configurations— can be applied to structural connectomes, cortical morphology, and transcriptomic gradients, suggesting the potential for a unified framework for quantifying asymmetry across multiple biological scales. Moreover, PDHC’s strong reliability and person specificity suggest utility as a trait-like endophenotype linking neural architecture, behavior, and genetic variation. Future studies could extend PDHC to task-dependent states and behavioral measures, clarifying its functional relevance for lateralized cognition. Applications to clinical populations with atypical lateralization provide an additional, mechanistically informative direction for future research.

In conclusion, PDHC provides a unified and quantitative framework for characterizing hemispheric functional asymmetry across individuals, developmental periods, and genetic contexts. By measuring divergence in connectivity patterns rather than directional biases, PDHC captures hemispheric differences at the level of large-scale network geometry and offers a principled basis for examining how bilateral coordination and hemisphere-specific specialization jointly shape human brain organization.

## Materials and Methods

### Datasets and study design

This study integrated four independent fMRI datasets to evaluate the reliability, developmental trajectory, and genetic basis of hemispheric functional organization: (1) the

Human Connectome Project (HCP) S1200 release (Van Essen et al., 2013); (2) an in-house Turner syndrome (TS) cohort; (3) the Developing Human Connectome Project (dHCP) (Fitzgibbon et al., 2020); and (4) the developmental component of the Chinese Color Nest Project (devCCNP) (Liu et al., 2021). The HCP dataset (n = 1,203) provided large-sample normative data for characterizing the main properties of PDHC, assessing its robustness across analytic setting, and testing short- and long-term reliability. The TS and HCP twin datasets were jointly analyzed to examine genetic contributions to PDHC. The dHCP (n = 505 neonates) and devCCNP (n = 196 children/adolescents) datasets were used to model age-related changes spanning late gestation through early adulthood. All data collection adhered to the ethical standards of the respective institutions, and informed consent was obtained from all participants or their legal guardians. Detailed demographic characteristics, MRI acquisition parameters, and quality-control procedures are provided in Supplementary Methods.

### fMRI preprocessing and hemispheric connectome construction

All datasets underwent standardized preprocessing consistent with the HCP minimal pipeline or equivalent procedures (Glasser et al., 2013), including slice-timing correction, motion correction, spatial normalization to template space, nuisance regression (global, white matter, and CSF signals), and temporal filtering (0.01–0.1 Hz for resting state). Full dataset-specific preprocessing pipelines are detailed in Supplementary Methods.

Two connectivity-based atlases were used to define hemispheric regions: the Atlas of Intrinsic Connectivity of Homotopic Areas (AICHA; 344 cortical + 40 subcortical regions) (Joliot et al., 2015) and the Human Brainnetome Atlas (BNA; 210 cortical + 36 subcortical regions)(Fan et al., 2016). For each subject, mean BOLD time series were extracted within each region, and intra-hemispheric functional connectivity (FC) matrices were computed as Pearson correlations between regional time series within each hemisphere. FC coefficients were Fisher z-transformed prior to statistical analysis. The resulting hemispheric FC matrices represented high-dimensional connectivity fingerprints for each hemisphere.

### Computation of the Pattern Dissimilarity of Hemispheric Functional Connectivity (PDHC)

For each subject, PDHC quantified the multivariate dissimilarity between left- and right-hemispheric FC matrices. Each hemispheric connectome was treated as a feature vector of all within-hemisphere connections (17,955 edges for AICHA; 7,503 for BNA). PDHC was defined as the Pearson correlation distance between hemispheric FC vectors:

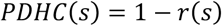

where *r(s)* is the Pearson correlation between the left and right hemispheric FC patterns of subject *s*. PDHC values range from 0 (perfectly symmetrical connectivity patterns) to 2 (completely dissimilar patterns). Lower PDHC indicates greater hemispheric similarity.

To establish population-level baselines, two inter-subject dissimilarity metrics were computed: ipsilateral-PDHC (between homologous hemispheres across individuals) and contralateral-PDHC (between non-homologous hemispheres across individuals). One-wayANOVAs and post hoc *t*-tests compared intra-subject PDHC to these baselines across sessions and atlases.

## Validation and reliability analyses

Reliability was assessed across multiple timescales. For short-term reproducibility, 960 HCP participants with both REST1 and REST2 scans (∼1 day apart) were analyzed. For long-term reproducibility, 42 subjects with retest data (interval = 1–11 months) were examined. Test–retest reliability was quantified by Pearson correlations and intraclass correlation coefficients (ICC). To control for potential genetic confounds, the same analyses were repeated in a subset of 93 unrelated HCP participants (HCP-U100).

Head-motion effects were evaluated by correlating mean framewise displacement (RMS) with PDHC values across participants in all datasets. To ensure robustness, analyses were repeated in low-motion neonatal subsets (mean RMS < 0.1 mm in neonates). Anatomical confounds were tested by correlating PDHC with whole-brain gray-matter asymmetry|(GMV_*lh*_ *– GMV*_*rh*_)/ (GMV_*lh*_ *+ GMV*_*rh*_)|. None of these potential confounds explained substantial variance in PDHC, confirming that PDHC reflects intrinsic hemispheric organization rather than motion or anatomical asymmetry.

### Network-level decomposition and enrichment analysis

To identify brain systems contributing most to hemispheric coupling, PDHC was decomposed into edge-wise standardized contributions. For each edge, a null distribution was derived from 1,000 random permutations disrupting the left–right pairing, and its contribution was then z-scored relative to this null distribution. Standardized edge contributions were mapped onto large-scale networks using two independent schemes: Mesulam’s hierarchical organization (Primary, Unimodal, Heteromodal, Paralimbic, Limbic/Subcortical)(Mesulam, 2000) and Yeo’s 7-network parcellation (VIS, SMN, DAN, VAN, LIN, FPN, DMN)(Yeo et al., 2011), extended with subcortical and divergent components.

To identify network pathways that disproportionately drive the strong interhemispheric coupling, we conducted an enrichment analysis. First, candidate edges were selected based on the largest-magnitude standardized contributions (top 3%, 5%, 10%, 15%, and 20%), all of which were negative. For each network pair, we computed the rich factor (RF), quantifying the normalized overrepresentation of candidate edges. Statistical significance was assessed via a null distribution generated from 5,000 permutations that shuffled edge contributions across the whole connectome, and p-values were FDR-corrected across network pairs (Benjamini & Hochberg, 1995).

### Developmental and genetic analyses

Age-related effects on PDHC were modeled across three developmental datasets (dHCP, devCCNP, and HCP). Linear mixed-effects (LME) models were fitted to longitudinaldatasets (dHCP, devCCNP) with postmenstrual age (PMA) or chronological age as fixed effects and subject identity as a random intercept and slope, controlling for sex, motion, and perinatal variables. For the cross-sectional HCP sample, multiple linear regression tested age effects on PDHC controlling for sex, handedness, and motion. Age associations were further validated using leave-one-out cross-validation, predicting PDHC from age without covariates.

Genetic influences were evaluated using two complementary approaches. First, heritability of PDHC was estimated in the HCP twin dataset using the Sequential Oligogenic Linkage Analysis Routines (SOLAR) with an AE variance-components model, controlling for age, age^2^, sex, and their interactions. Second, X-chromosome dosage effects were assessed in Turner syndrome (TS) patients (45,XO) versus age-matched typically developing girls using two-sample t-tests controlling for scan age.

These analyses jointly provided converging evidence for genetic contributions to hemispheric pattern-level asymmetry.

## Supporting information

DataS1

## Final note

Comprehensive acquisition parameters, preprocessing pipelines, and supplementary analyses are provided in Supplementary Methods.

## Acknowledgments

This work is supported by the National Natural Science Foundation of China (Nos. T2325006, 82021004), STI 2030-Major Projects (Nos.2021ZD0201701,2021ZD0200500); the Fundamental Research Funds for the Central Universities (No. 2233200020).

## Author contributions

Conceptualization: G.G, C. Z, Q.B

Methodology: Q.B, C. Z, X-N. Z, S.P

Investigation: Q.B, C. Z

Visualization: Q.B, C.

ZSupervision: G.G, B.S

Writing—original draft: Q.B, C. Z

Writing—review & editing: G.G, Q.B

## Competing interests

Authors declare that they have no competing interests.

## Data and materials availability

All data are available at request.

## Notes

### Competing Interest Statement

The authors have declared no competing interest.

