## Supplementary Materials for "A multivariate pattern metric of individualized hemispheric functional asymmetry"

**Supplementary Methods**

- **Datasets**

**Human Connectome Project (HCP) dataset**

The S1200 release of the Human Connectome Project (HCP) was adopted for primary analyses of PDHC characteristics and methodological evaluation. The dataset includes multimodal MRI data from over 1,200 healthy young adults (ages 22–35). All participants provided written informed consent under protocols approved by the Institutional Review Board of Washington University.

**MRI acquisition.** Functional MRI data were collected using a customized 3T Siemens Connectome Skyra scanner with a 32-channel head coil. Two resting-state fMRI (rs-fMRI) sessions (REST1, REST2) were acquired over two days. Imaging parameters were identical across sessions: multiband EPI sequence (factor 8); TR = 720 ms; TE = 33.1 ms; flip angle = 52°; bandwidth = 2290 Hz/pixel; FOV = 208 × 180 mm^2^; 72 slices; voxel size = 2 mm isotropic; 1200 time points per rs-fMRI run; with both left–right (LR) and right–left (RL) phase-encoding directions.

**fMRI preprocessing.** The HCP minimal preprocessing pipeline was applied, including gradient distortion correction, motion and EPI distortion correction, registration to T1-weighted images and MNI space, global intensity normalization, and brain extraction (Glasser et al. 2013). rs-fMRI data were further denoised using FIX (ICA-based Xnoisifier) to remove structured noise (Smith et al. 2013; Salimi-Khorshidi et al. 2014). Additional preprocessing in GRETNA (Wang et al. 2015) included linear detrending, nuisance regression (global, white-matter, and cerebrospinal-fluid signals), and temporal bandpass filtering (0.01–0.1 Hz).

**Quality control.** Following HCP recommendations, rs-fMRI scans labeled with QC codes A were excluded. Participants missing either LR or RL runs were removed. REST1 data from 1,033 subjects (470 males) and REST2 data from 968 subjects (446 males) were retained.

**HCP retest dataset**

A subset of 46 HCP participants underwent a second identical imaging session (interval: 1–11 months). After QC, 42 participants for REST1 and 41 for REST2 were included. All data were preprocessed using the same pipeline as the main HCP pipeline. These data were used to evaluate long-term test–retest reliability.

**In-house Turner Syndrome (TS) dataset**

An in-house Turner syndrome (TS) dataset was used to investigate the X-chromosomal contribution to PDHC. The sample included 22 girls with confirmed nonmosaic 45,XO karyotype (ages 10.2–17.6 years) and 21 age-matched controls (ages 9.9–17.5 years). Cytogenetic diagnosis was verified using peripheral blood. Participants had no history of neurological or psychiatric disorders. The study was approved by the Research Ethics Committee of Beijing Normal University, and informed consent was obtained from legal guardians.

**MRI acquisition.** Scanning was performed on a 3T Siemens Tim Trio system (Beijing Normal University) using a 12-channel head coil.

rs-fMRI: TR = 2000 ms; TE = 30 ms; flip angle = 90°; 33 slices; slice thickness/gap = 3.5/0.7 mm; voxel size = 3.1 × 3.1 mm^2^; 200 time points.

T1-weighted: MPRAGE, TR = 2530 ms; TE = 3.39 ms; TI = 1100 ms; 144 slices; voxel size = 1 × 1 × 1.33 mm^3^.

**Preprocessing.** All rs-fMRI images were preprocessed using the GRETNA toolbox. The first 10 volumes were discarded. Images underwent slice-timing correction, motion realignment, T1 co-registration, DARTEL normalization to MNI space, and nuisance regression (Friston 24-parameter motion model, global, WM, and CSF signals). Temporal bandpass filtering (0.01–0.1 Hz) and linear detrending were applied.

**Quality control.** Subjects exceeding 2 mm or 2° motion were excluded. The final sample comprised 22 TS and 18 controls (mean ages 13.8 ± 2.4 and 14.2 ± 2.3 years, respectively).

**Developing Human Connectome Project (dHCP) dataset**

The dHCP dataset was used to investigate early-life PDHC development (https://developingconnectome.org). Neonatal MRI was approved by the UK National Research Ethics Committee, with informed consent from parents. We used the first and second data releases (totaling 538 rs-fMRI scans, 480 neonates; PMA = 24–45 weeks).

**MRI acquisition.** Scans were obtained at Evelina London Children’s Hospital on a 3T Philips Achieva system using a dedicated neonatal head coil. rs-fMRI: multiband EPI, MB = 9; TE = 38 ms; TR = 392 ms; voxel size = 2.15 mm isotropic; 2300 volumes per run. Most infants were imaged during natural sleep.

**Preprocessing.** The dHCP pipeline (Hughes et al. 2017) performed distortion and motion correction, multimodal registration, ICA-based denoising, and automated QC. Additional steps (using the GRETNA toolbox) included regression of global/WM/CSF signals and temporal bandpass filtering (0.01–0.1 Hz).

**Quality control.** Of 538 scans, 512 passed automated QC. Following radiological review, 463 scans from 430 infants (mean birth PMA 37.7 ± 4.0 weeks; 244 males) were retained. Among them, 33 preterm infants were scanned twice (once preterm and once at term-equivalent age).

**Developing Chinese Color Nest Project (devCCNP) dataset**

The devCCNP dataset was used to examine PDHC maturation from childhood through adolescence (Liu et al. 2021). It included 198 participants (ages 6–18) scanned longitudinally up to three times (mean interval = 1.3 ± 0.2 years). The study was approved by the Research Ethics Committee of Southwest University, Chongqing.

**MRI acquisition.** Scans were acquired on a 3T Siemens Tim Trio scanner using a 12-channel head coil.

rs-fMRI: TR = 2500 ms; TE = 30 ms; flip angle = 80°; voxel size = 3.0 mm isotropic; 184 time points per run.

T1-weighted: TR = 2600 ms; TE = 3.02 ms; TI = 900 ms; voxel size = 1 × 1 × 1 mm^3^.

**Preprocessing.** Preprocessing was conducted using the Connectome Computation System (CCS) (Xu et al. 2015). Steps included discarding the first 4 initial volumes, slice-timing and motion correction, ICA-AROMA motion artifact removal, regression of global/WM/CSF signals, removal of linear trends, spatial smoothing (FWHM = 6 mm), boundary-based registration (Greve and Fischl 2009), and normalization via DARTEL to MNI space. Given ICA-AROMA’s efficacy, no bandpass filtering was applied (Pruim et al. 2015).

**Quality control.** Scans lacking full brain coverage or missing one of two runs were excluded. A total of 419 rs-fMRI scans from 196 participants (91 males) were included.

- **Functional connectome reconstruction**

Preprocessed fMRI data were aligned to either MNI (HCP and devCCNP) or 40-week neonatal template (dHCP) space. Mean BOLD time series were extracted from each region of interest (ROI) within each hemisphere. Intra-hemispheric functional connectivity (FC) was computed as the Pearson correlation between ROI time series, and correlations were Fisher z-transformed. For the HCP dataset, FC matrices from LR and RL runs were averaged to mitigate phase-encoding effects. Each hemispheric connectome thus represented a square FC matrix, later flattened into a 1D vector of within-hemispheric edges (17,955 for AICHA; 7,503 for BNA).

- **Computation of PHDC**

The Pattern Dissimilarity of Hemispheric Functional Connectivity (PDHC) quantified the multivariate dissimilarity between left and right hemispheric FC patterns for each subject:
 $PDHC\left( s \right)=1-r\left( s \right)$ (S1)where *r(s)* is the Pearson correlation between Fisher z-transformed FC vectors of the left and right hemispheres for subject *s*. PDHC ranges from 0 (perfect similarity) to 2 (complete dissimilarity).

Two inter-subject baselines were computed: (1) Ipsilateral PDHC, the correlation distance between homologous hemispheres across subjects, and (2) Contralateral PDHC, the distance between non-homologous hemispheres across subjects. One-way ANOVAs followed by t-tests compared intra-subject PDHC with these baseline values.

Subject-specificity was further tested via a subject identification analysis adapted from Finn et al. (Finn et al. 2015). For each participant, the FC pattern from one hemisphere was used as a target and correlated with all hemispheres in the database. Identification was successful if the intra-subject PDHC was the smallest distance among all pairings. Analyses were repeated for REST1, REST2, and cross-session combinations. Null distributions were estimated from 1,000 random permutations of subject label.

- **Network-level decomposition and enrichment**

**Edgewise decomposition**

Edgewise contributions to PDHC were computed as:
 $\varphi\left( s,e \right)=-\frac{1}{n}\cdot Z_{lh}\left( s,e \right)\cdot Z_{rh}\left( s,e \right)$	(S2)
where *Z_lh_* and *Z_rh_* are z-scored FC values for edge *e* in the left and right hemispheres, and n is the total number of intra-hemispheric edges. Group-level edgewise contributions were averaged across subjects.

**Standardization and null model**

A non-parametric null distribution was generated for each edge by shuffling the right-hemisphere subject labels 1,000 times. The observed group-level edge contribution was converted into a z-score relative to this null distribution, yielding the standardized edge contribution. Negative values indicate edges that reduce hemispheric dissimilarity, whereas positive values indicate edges that promote dissimilarity.

**Functional network mapping**

Standardized edge contributions were mapped to large-scale networks using two organizational schemes: (1) Mesulam’s hierarchical organization (Primary, Unimodal, Heteromodal, Paralimbic, Limbic/Subcortical) (Mesulam 2000) and (2) Yeo’s 7-network parcellation (Yeo et al. 2011), expanded to nine components (VIS, SMN, DAN, VAN, LIN, FPN, DMN, SUB, DIV).

**Enrichment analysis**

Edges with the largest absolute standardized contributions were selected (top 3–20%). The rich factor (RF) quantified the normalized proportion of selected edges within a given network. Significance was tested using 5,000 permutations, generating a null distribution of RF values separately for each network under random assignment. P-values were corrected for multiple comparisons (Benjamini and Hochberg 1995).

- **Developmental and genetic analysis**

**Developmental modeling**

Age effects on PDHC were analyzed using the *lme4* (R 4.3.3):
- dHCP: linear mixed-effects model; PMA as fixed effect; subject identity as random intercept and slope; covariates = sex, delivery type, birth weight, head circumference, and motion (RMS displacement).
- devCCNP: linear mixed-effects model; age as fixed effect; subject identity as random intercept and slope; covariates = sex, total brain volume, and motion (RMS displacement).
- HCP: linear model; covariates = sex, handedness, motion (RMS displacement).

Interaction terms (age × sex) were tested; if nonsignificant, only main effects were reported. Leave-one-out cross-validation (LOOCV) using models without covariates estimated PDHC predictability from age.

**Heritability estimation**

Genetic influences were examined using HCP twin data. Heritability (h^2^) was estimated with the AE variance-components model in SOLAR (Almasy and Blangero 1998). PDHC values were inverse-normal transformed before analysis. Covariates included age, age^2^, sex, and their interactions.

**X-chromosome dosage effect**

To assess X-chromosome dosage contributions, PDHC was compared between Turner syndrome (45,XO) and control girls using two-sample t-tests within a GLM framework that controlled for age.

- **Validation analysis**

Long-term reproducibility was assessed using intraclass correlation coefficients (ICC) between first and follow-up scans (1–11 months). Motion effects were evaluated via mixed-effects models correlating PDHC with mean RMS displacement. Analyses for dHCP were repeated in a low-motion subset (mean RMS < 0.1 mm). Anatomical confounds were tested by correlating PDHC with absolute cortical GMV asymmetry |(GMV_lh_ - GMV_rh_)| / (GMV_lh_ + GMV_rh_).

To address potential genetic confounds, analyses were replicated in the HCP-U100 unrelated subset, confirming robustness across parcellations and sessions.

**Supplementary Figures**

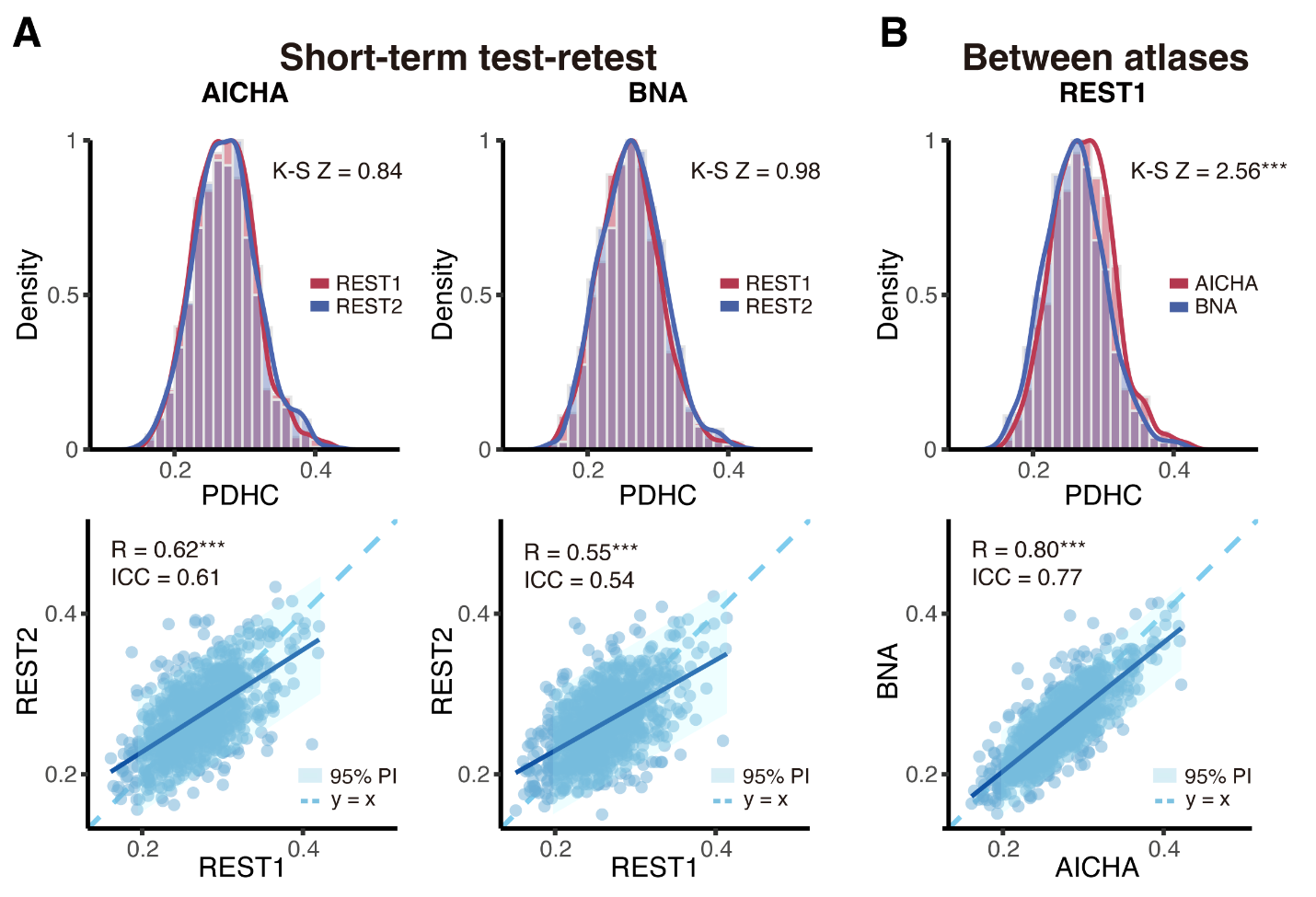

**Fig. S1. Reproducibility and cross-atlas robustness of PDHC without global signal regression (GSR).** (A) Short-term (one-day) test–retest reproducibility of PDHC across REST1 and REST2 sessions after excluding global signal regression. PDHC distributions largely overlapped, and inter-session correlations remained strong (ICC = 0.54–0.61). (B) Cross-atlas correspondence between AICHA and BNA under identical preprocessing condition. Despite minor distributional differences (as indicated by K–S tests significant), inter-atlas correlations were high (ICC = 0.77), confirming the robustness of PDHC to atlas selection and preprocessing choices. Abbreviations: PDHC, Pattern Dissimilarity of Hemispheric Functional Connectivity; FC, functional connectivity; AICHA, Atlas of Intrinsic Connectivity of Homotopic Areas; BNA, Brainnetome Atlas; PI, prediction interval.

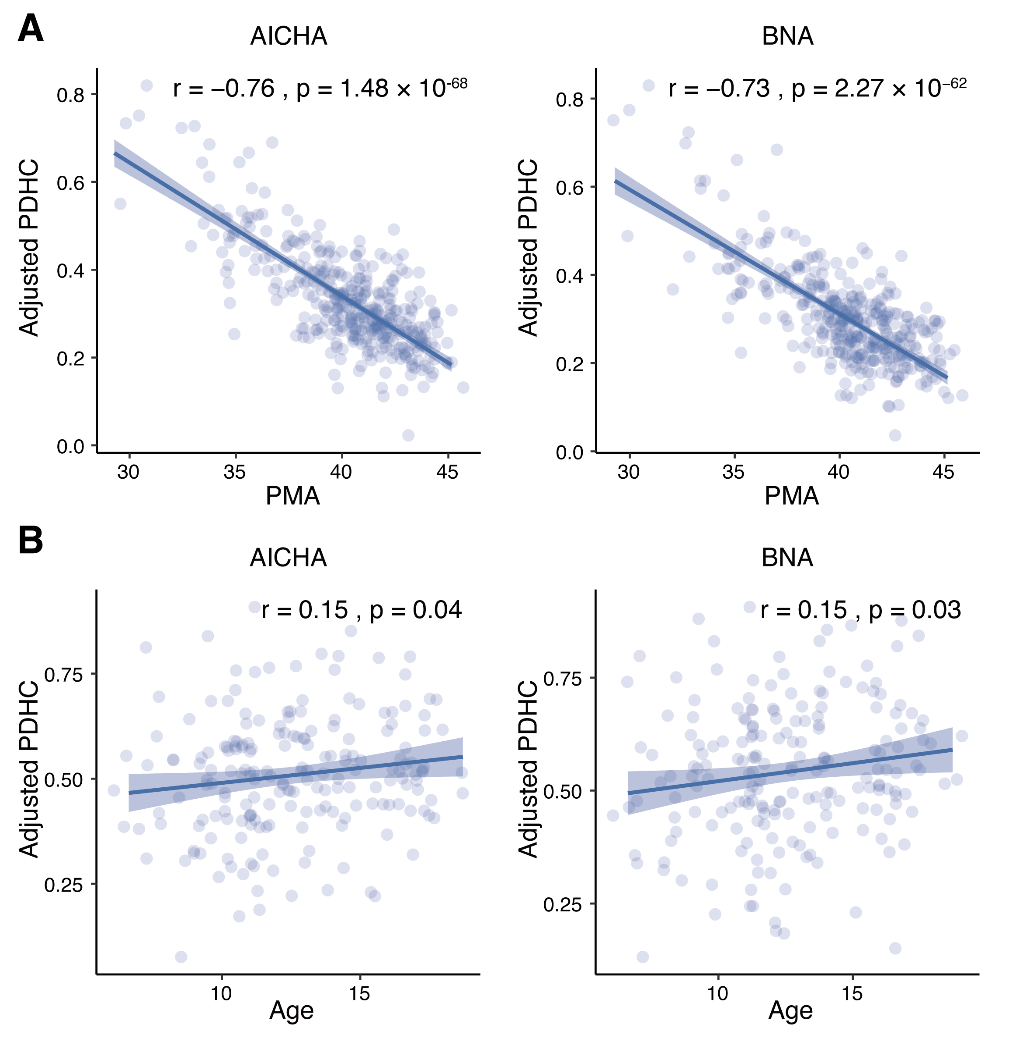

**Fig. S2. Validation of developmental effects of PDHC using independent-session data.** (A) In the dHCP neonate cohort, PDHC decreased markedly with postmenstrual age (PMA) for both AICHA and BNA atlases, confirming rapid early differentiation of hemispheric organization. These analyses were restricted to each infant’s first available scan to avoid potential repeated-measure dependencies. (B) In the devCCNP cohort, PDHC increased modestly with age from childhood to adolescence. Similarly, only the first longitudinal session of each participant was used to prevent within-subject repetition effects. Linear models controlled for relevant covariates: delivery type, birth weight, sex, head circumference, and motion for the dHCP dataset; and sex, total brain size, and mean RMS head motion for the devCCNP dataset. Abbreviations: PDHC, Pattern Dissimilarity of Hemispheric Functional Connectivity; PMA, postmenstrual age; AICHA, Atlas of Intrinsic Connectivity of Homotopic Areas; BNA, Brainnetome Atlas; dHCP, developing Human Connectome Project; devCCNP, developmental Chinese Color Nest Project.

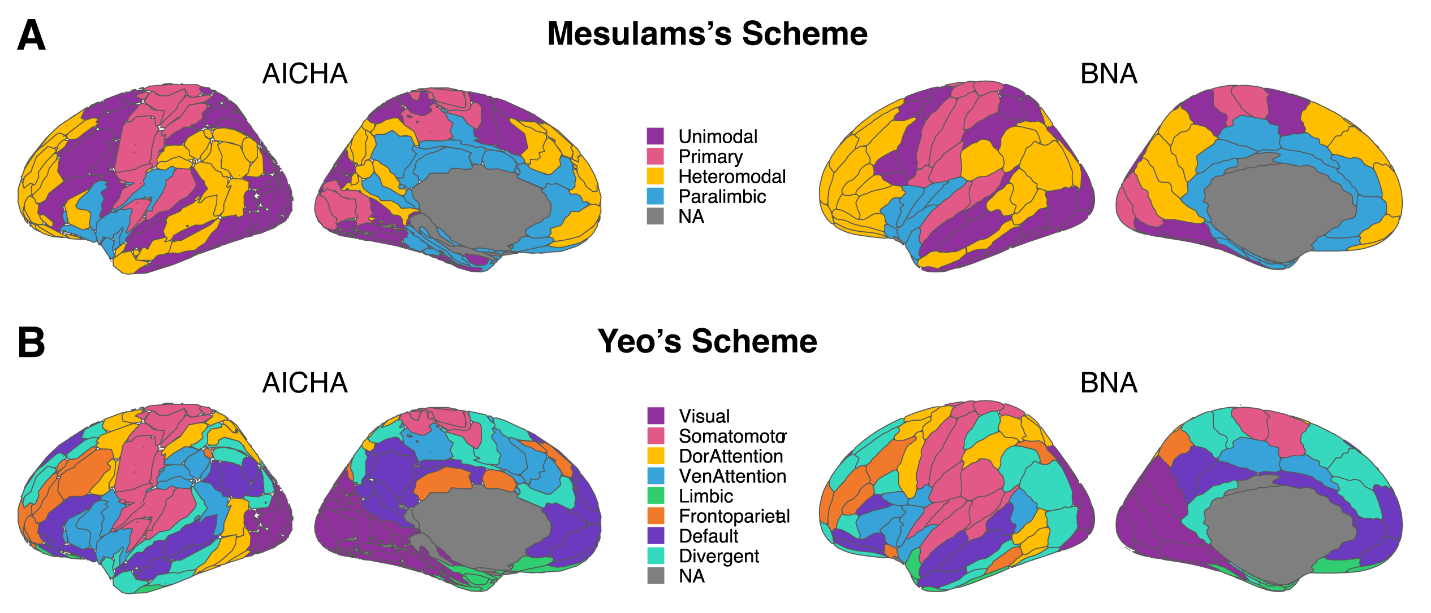

**Fig. S3.** **Region-to-network mappings for AICHA and BNA atlases.** (A) Mapping of AICHA and BNA atlas regions onto Mesulam’s hierarchical classification, including Primary, Unimodal, Heteromodal, Paralimbic, and Limbic/Subcortical levels. These mappings provide the organizational basis for system-level decomposition of PDHC edge contributions. (B) Mapping of the same atlases onto Yeo’s 7-network framework, comprising Visual (VIS), Somatomotor (SMN), Dorsal Attention (DAN), Ventral Attention (VAN), Limbic (LIN), Frontoparietal (FPN), Default Mode (DMN), and Subcortical (Subc) systems. Full mapping tables are provided in Supplementary Data S1, with network-level region counts summarized in Supplementary Table S1. Abbreviations: AICHA, Atlas of Intrinsic Connectivity of Homotopic Areas; BNA, Brainnetome Atlas.

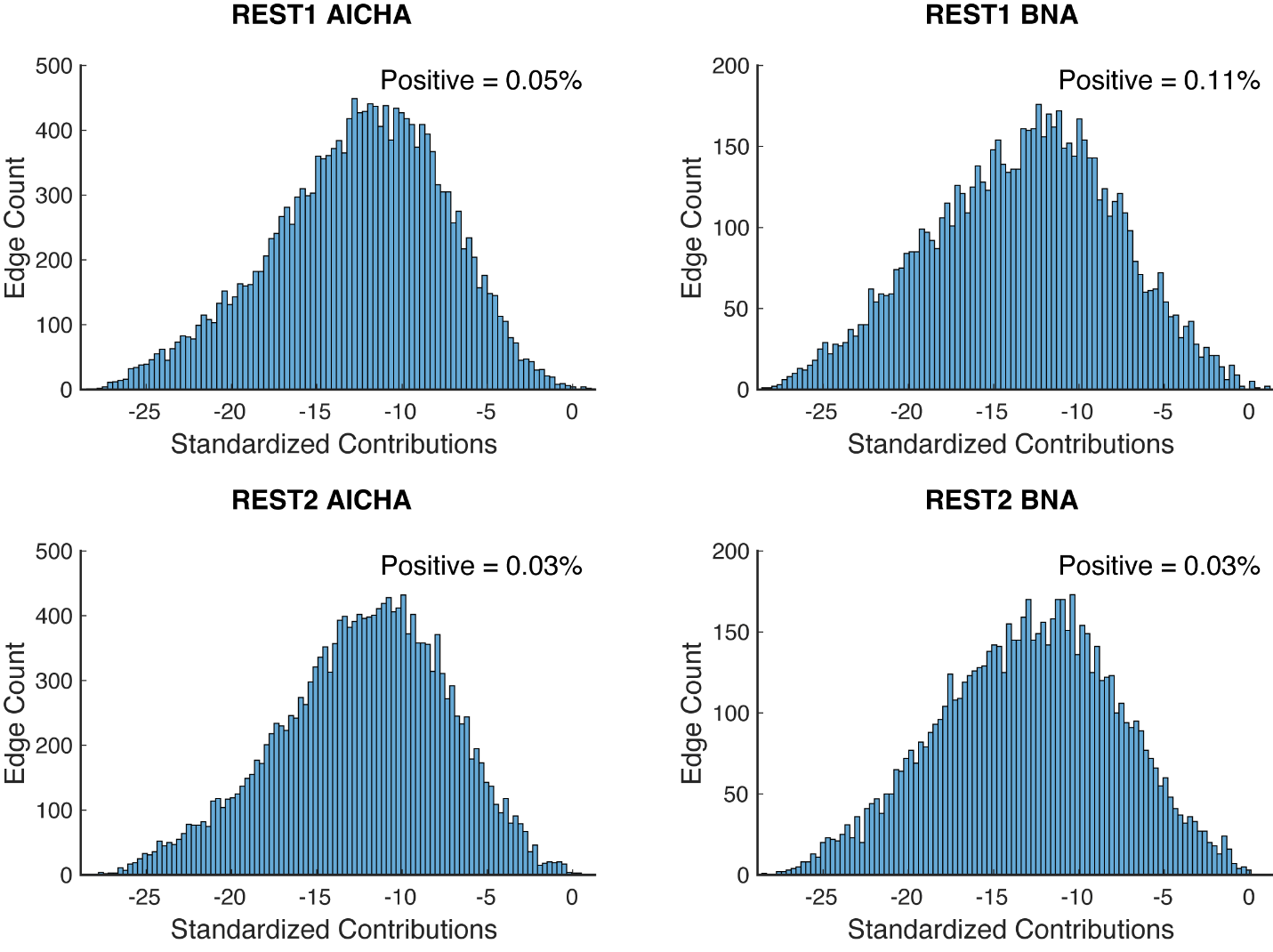

**Fig. S4. Distribution of edge-wise standardized contributions to PDHC.** Histograms show the z-scored edge-wise contributions for REST1 and REST2 sessions using the AICHA (17,955 edges) and BNA (7,503 edges) atlases. Negative z-scores represent edges that reduce hemispheric dissimilarity (lower PDHC), whereas positive z-scores indicate edges increasing hemispheric dissimilarity. Across both atlases and sessions, the vast majority of edges exhibited negative standardized contributions. Abbreviations: PDHC, Pattern Dissimilarity of Hemispheric Functional Connectivity; AICHA, Atlas of Intrinsic Connectivity of Homotopic Areas; BNA, Brainnetome Atlas.

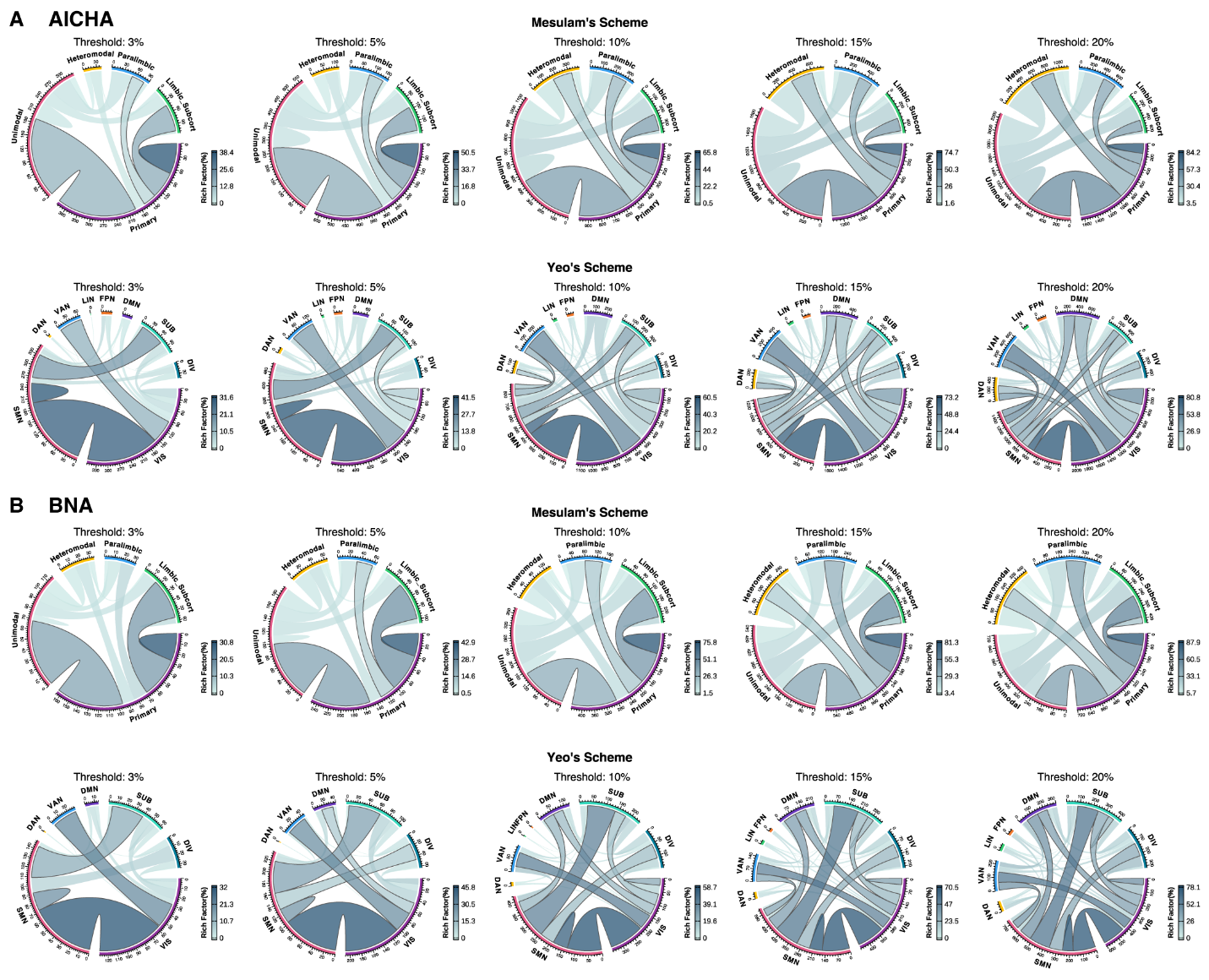

**Fig. S5. Network-level enrichment of hemispheric coupling across edge-selection thresholds.** Chord plots show enrichment results from REST1 data across thresholds of 3%, 5%, 10%, 15%, and 20%, generated using the R *circlize* package. (A) AICHA and (B) BNA atlas results are displayed separately. Line color represents the rich factor (RF), line width corresponds to the number of supra-threshold edges, and bold borders denote significance after FDR correction (p_FDR_ < 0.05). Across all thresholds, enrichment consistently concentrated within lower-order systems—particularly Mesulam’s Primary network and Yeo’s Visual and Somatomotor networks—demonstrating the dominant role of conserved sensorimotor architecture in reducing hemispheric dissimilarity. Abbreviations: AICHA, Atlas of Intrinsic Connectivity of Homotopic Areas; BNA, Brainnetome Atlas; RF, rich factor; FDR, false discovery rate.

**Supplementary Tables**

**Table S1. Demographic information.**

|  | Number(M/F) | Scan age  (years/weeks PMA) | Birth age  (weeks PMA) |
| --- | --- | --- | --- |
| **HCP Dataset** |  |  |  |
| Young adults (REST1) | 1033(470/563) | 28.78 ± 3.70 | - |
| Young adults (REST2) | 968(446/522) | 28.71 ± 3.71 | - |
| **HCP Retest Dataset** |  |  |  |
| Young adults (REST1) | 42(12/30) | 30.45 ± 3.35 | - |
| Young adults (REST2) | 41(13/28) | 30.24 ± 3.43 | - |
| **TS Dataset** |  |  |  |
| Healthy control | 18(-/18) | 14.2 ± 2.3 | - |
| Turner Syndrome | 22(-/22) | 13.8 ± 2.4 | - |
| **dHCP Dataset** |  |  |  |
| Neonates  Total scans | 430(244/186)  463(268/195) | 40.21 ± 2.85 | 37.7 ± 4.05 |
| *Including* |  |  |  |
| Term-born babies | 397(220/177) | 40.63 ± 2.39 | 38.67 ± 3.35 |
| *Preterm-born babies | 33(24/9) | 1^st^: 34.19 ± 1.98  2^nd^: 41.19 ± 1.46 | 31.94 ± 2.98 |
| **devCCNP Dataset** |  |  |  |
| Children/Adolescents  Total scans | 196(91/105)  419(199/220) | 12.21 ± 2.93 | - |

*The preterm-born babies were scanned twice. PMA, postmenstrual age. dHCP, developing human connectome project. devCCNP, developing part of the Chinese Color Nest Project.

**Table S2. Summary of region assignments in AICHA and BNA atlases.**

| **Network** | **#AICHA Regions** | **#BNA Regions** |
| --- | --- | --- |
| Mesulam: Primary | 20 | 14 |
| Mesulam: Unimodal | 67 | 34 |
| Mesulam: Heteromodal | 56 | 35 |
| Mesulam: Paralimbic | 27 | 22 |
| Mesulam: Limbic/Subcortical | 20 | 18 |
| Yeo: Visual (VIS) | 35 | 15 |
| Yeo: Somatomotor (SMN) | 19 | 15 |
| Yeo: Dorsal Attention (DAN) | 16 | 12 |
| Yeo: Ventral Attention (VAN) | 15 | 9 |
| Yeo: Limbic (LIN) | 14 | 11 |
| Yeo: Frontoparietal (FPN) | 15 | 8 |
| Yeo: Default (DMN) | 30 | 14 |
| Yeo: Subcortical (SUB) | 20 | 18 |
| Yeo: Divergent (DIV) | 26 | 21 |
| **Total** | **190** | **123** |
| This table summarizes the number of hemispheric regions assigned to each network for AICHA (190 regions) and BNA (123 regions) atlases. Full mappings are listed in Supplementary Data S1. | | |

**Table S3. (separate file)**

Enrichment analysis results using Mesulam’s scheme at 3% threshold for AICHA and BNA atlases.

**Table S4. (separate file)**

Enrichment analysis using Yeo’s scheme at 3% threshold for AICHA and BNA atlases.

**Data S1. (separate file)**

Type or paste caption here.
